# Reproducibility and cross-dataset concordance of drug-induced transcriptional responses

**DOI:** 10.64898/2026.11.11.765551

**Authors:** Jinming Bai, Sharon Prince, Geoff S. Nitschke

**Affiliations:** Department of Human Biology, Faculty of Health Sciences, University of Cape Town, Cape Town 7925, South Africa; Department of Computer Science, School of Information Technology, Faculty of Science, University of Cape Town, Cape Town 7701, South Africa

## Abstract

Drug-perturbation transcriptomic resources are expanding, but the reproducibility and cross-source consistency of the responses they measure remain critical to their use. We compared six resources using independent replicates, matched-drug analyses, and response decomposition. High agreement in treated expression did not translate into equally repeatable drug-induced changes. Cross-source response concordance was generally limited, including under matched cell and treatment conditions, although some comparisons retained drug-specific information. In Tahoe, a control-expression-only baseline left substantial drug-specific variation unresolved. Measured drug main effects-the average responses across cell lines-provided information for predicting cell-specific interactions, with this relationship retained in a cross-plate validation. Exploratory analyses of structure, target, and external-response representations identified modest signals in selected pipelines. Together, these analyses characterize the reliability and transferability of current perturbation resources and identify prediction of drug main effects as a key bottleneck in the framework examined, motivating shared reference measurements and better recovery of drug-specific responses.

**Highlights:**

- Expression similarity does not establish drug-response reproducibility
- Matched drug responses often show limited agreement across data sources
- Drug-specific components account for most error in a fixed control-expression baseline
- Measured drug effects predict interactions across a tested plate pair

## Introduction

Transcriptional responses to chemical perturbations are used to investigate drug mechanisms and develop models of cellular responses. By measuring many compounds across cell lines and experimental conditions, resources such as Tahoe-100M, L1000, DRUG-seq, sciPlex3, CIGS, and MIX-seq create opportunities to compare response patterns and test predictions beyond individual experiments.[1–6] Drugs shared between resources could connect these measurements and help assess whether a response generalizes beyond its original experiment. These opportunities depend on what the measurements capture. The resources differ in gene coverage, cellular composition, dose and treatment duration, control design, and data processing, so a shared compound name does not by itself establish a comparable response.

Three properties are therefore important to distinguish. Repeatability asks whether independently measured responses agree within a source under matched conditions. Cross-source concordance asks whether matched perturbations agree across resources, conditional on the cells, doses, times, and genes available for comparison. Predictive utility asks whether information improves a specified prediction for held-out observations. None of these properties alone establishes the others. High treated-expression correlation can be dominated by persistent cellular background, whereas subtracting controls changes both the signal and its uncertainty. Low response-vector concordance can coexist with information that distinguishes one drug from alternatives. Conversely, a reproducible association with one response component may fail to improve the complete response.

Model evaluation makes this distinction particularly consequential. A predictor can recover shared cellular or experimental variation while capturing little information that distinguishes drugs exposed to the same background. Systema demonstrated a related problem in genetic-perturbation prediction and introduced perturbation-specific evaluation references.[7] For chemical perturbations, this motivates a complementary measurement-level question: which parts of the observed response repeat, agree across sources, and remain unresolved by a model without drug information? Drug main effects describe mean responses across cells relative to the overall mean; interactions are cell-dependent departures from the additive cell and drug effects.[8] Addressing this question requires attention to experimental support as well as scores. Correlations computed on different gene axes or drug sets need not be directly comparable, and an interaction residual includes measurement error as well as potentially informative biology.

Here, we examine these questions through a coordinated audit of six transcriptomic resources and an illustrative, fixed Tahoe predictor. We compare treated expression and control-subtracted responses on native and common gene axes, distinguish source-wide descriptions from matched cross-source comparisons, and relate vector agreement to drug discrimination. We then decompose observed responses to identify how experimental design constrains component interpretation. In Tahoe, a control-expression baseline provides an explicit reference for locating drug-specific residual information, while measured drug main effects allow us to test their relationship with interactions across one independently measured plate pair. The aim is to clarify the evidence and remaining bottlenecks for response transfer and prediction, while keeping measurement agreement, component-level associations, and complete-response utility distinct.

## Results

### Drug-response repeatability is limited in most datasets

We first tested whether repeating the same drug treatment yielded consistent changes in gene expression. Across Tahoe, L1000 L3, DRUG-seq, CIGS, sciPlex3, and MIX-seq, we compared normalized treated expression with drug responses defined as treated-minus-control expression in independent replicates matched for drug, cell line, dose, and treatment duration. We included all eligible repeats in the analyzed public releases and evaluated each source’s native measured genes alongside a fixed set of 869 shared genes.

Treated expression was highly correlated between repeats, whereas drug-response correlations were substantially lower on both gene sets (Figures 1A and 1B). On the shared gene set, Tahoe retained moderate response repeatability, while L1000 L3, DRUG-seq, CIGS, and sciPlex3 showed weak average response correlations. MIX-seq had a higher response correlation than several other sources, but only one drug had qualifying independent repeats, precluding a dataset-wide interpretation. High agreement in treated expression therefore did not establish comparable repeatability of the changes induced by a drug. Response repeatability needs to be assessed directly from expression changes rather than inferred from the similarity of treated expression profiles.

The estimates also depended on which genes were evaluated. With replicate pairs held fixed, native and shared gene sets gave different response correlations, and the direction of the difference varied across sources: Tahoe showed higher correlation on its native gene set, whereas DRUG-seq showed higher correlation on the shared set (Figure 1B). The shared genes provide a consistent evaluation space, while the native genes retain information about each source’s own measurement coverage. Neither estimate should be interpreted independently of the gene set used.

Processing choices further affected repeatability. With genes and replicate pairs held fixed, increasing the pseudocount before log transformation raised response correlations in DRUG-seq, CIGS, and sciPlex3 (Figure S1A). The transformation compresses differences at low expression and may attenuate both noise and weak biological responses, so a higher correlation alone does not establish greater measurement fidelity. Together, these analyses show that drug-induced changes were only partly reproducible under the conditions examined. They do not, by themselves, distinguish technical noise from biological variability or establish a fixed limit on predictability.

**Figure 1.**
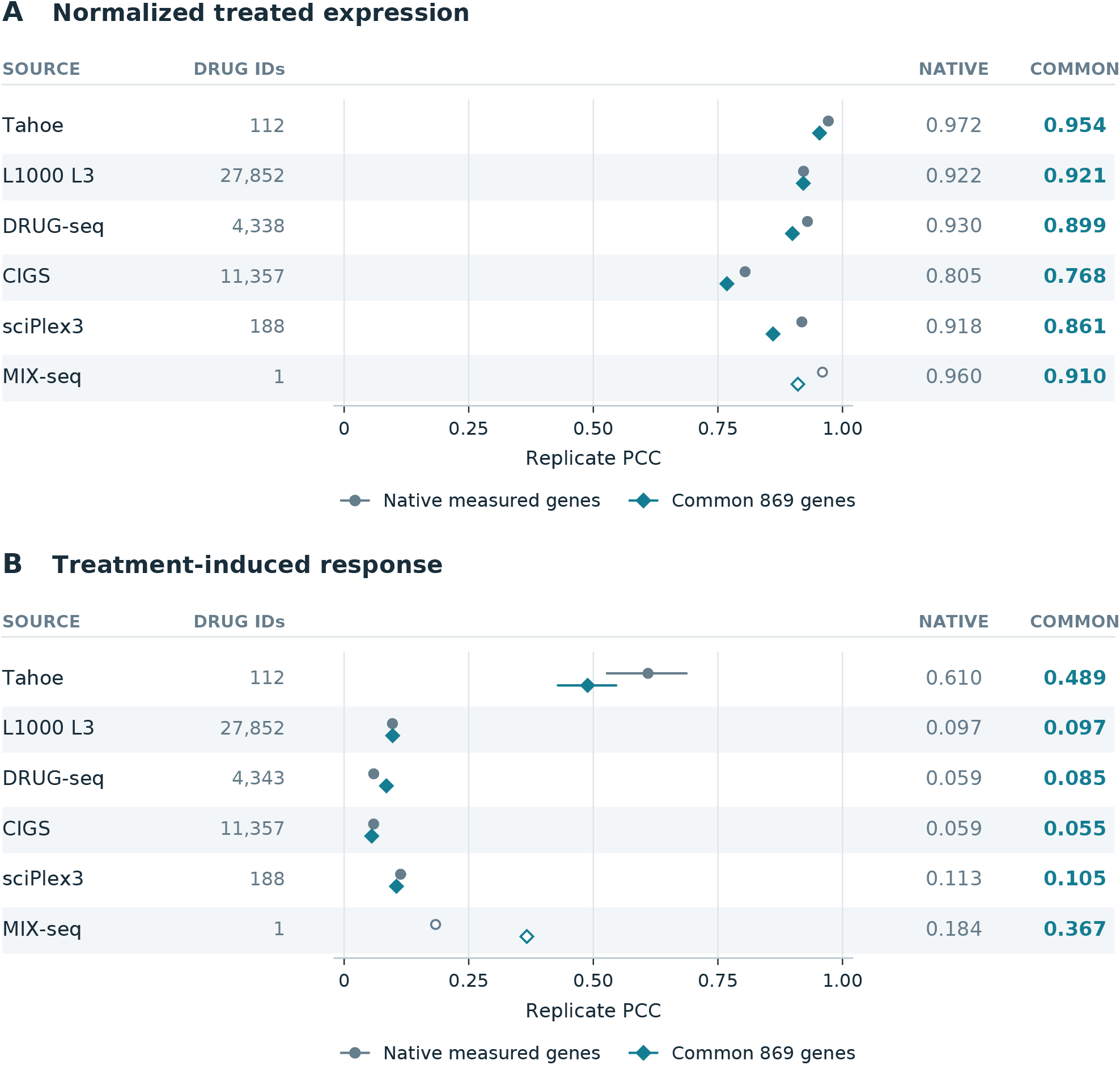
Repeatability of treated expression and drug responses. A, correlations between independent replicates of normalized drug-treated expression. B, correlations between independent drug responses, defined as treated expression minus matched control expression. Each source is evaluated on its native measured genes and 869 common genes, using identical replicate pairs across gene sets within each endpoint. n denotes contributing drug IDs. Points show mean PCC and horizontal lines show 95% bootstrap intervals; open points indicate fewer than five drugs. MIX-seq has one qualifying repeat drug and no drug-level interval.

### Cross-dataset drug responses show limited concordance but retain some drug-specific information

We next asked whether the same drug induced similar expression changes in different datasets. Within the shared gene space, we compared responses at matched treatment durations and nearby doses, considering both responses averaged across each source’s available cell lines and responses measured in the same cell line. These comparisons assess agreement in average drug responses and in responses under a more closely matched cellular background, respectively.

Tahoe showed the highest average response repeatability among datasets supported by multiple repeat-eligible drugs, providing a useful starting point for cross-source comparisons. Nevertheless, its cell-averaged drug responses generally showed low correlations with those in external datasets (Figure 2A). Restricting comparisons to the same cell line did not consistently yield strong agreement (Figure 2B). For example, Tahoe and L1000 L3 had an average same-cell correlation of approximately 0.014 across 189 matched drugs. Matching cell identity and nearby doses was therefore insufficient to ensure that the two sources recorded similar drug-response patterns. Because the cell-averaged and same-cell analyses included different drug and cell supports, their numerical differences cannot be attributed directly to cellular background.

To examine whether unequal comparison supports accounted for the low concordance, we restricted the analysis to conditions with independent repeats in both sources and calculated within-source repeatability and cross-source agreement on identical support. In all five comparisons containing at least five drugs, the cross-source point estimate was lower than both within-source repeatability estimates (Figure S1B). Among the same 49 drugs shared by Tahoe and L1000 L3, cross-source correlation was approximately 0.010, compared with within-source correlations of 0.471 and 0.106, respectively. Thus, limited cross-source agreement persisted after aligning the observations used for comparison. These differences do not, however, isolate the contributions of measurement platform, experimental implementation, or cell state.

The pattern extended to comparisons among external datasets. L1000 L3 and CIGS shared 2,505 matched drugs, yet their average cell-averaged response correlation was only 0.046 (Figure 2A). Extensive drug overlap therefore did not necessarily imply consistent response patterns. Some combinations showed higher concordance, including L1000 L3 and MIX-seq, but this comparison involved only 11 drugs. Cross-source agreement varied among dataset pairs and remained conditional on the available drug support.

Low response correlation did not imply a complete absence of shared drug-specific information. Using Tahoe responses to query external datasets, we compared same-drug matches with different-drug matches within the same experimental background. We also calculated retrieval AUC, the probability that the correct drug received a higher similarity score than an alternative. L1000 L3, CIGS, and sciPlex3 retained a same-drug matching advantage and retrieval AUCs above chance (Figures 2C and 2D). The Tahoe-L1000 L3 AUC was approximately 0.55, indicating modest discrimination rather than reliable drug identification. The DRUG-seq interval included chance, while MIX-seq contributed only three eligible query drugs and did not support a general conclusion.

These results show that some drug-specific information is preserved across datasets even when their complete response vectors agree poorly. Distinguishing the correct drug from alternatives does not establish that a response measured in one source can accurately represent the response recorded in another.

**Figure 2.**
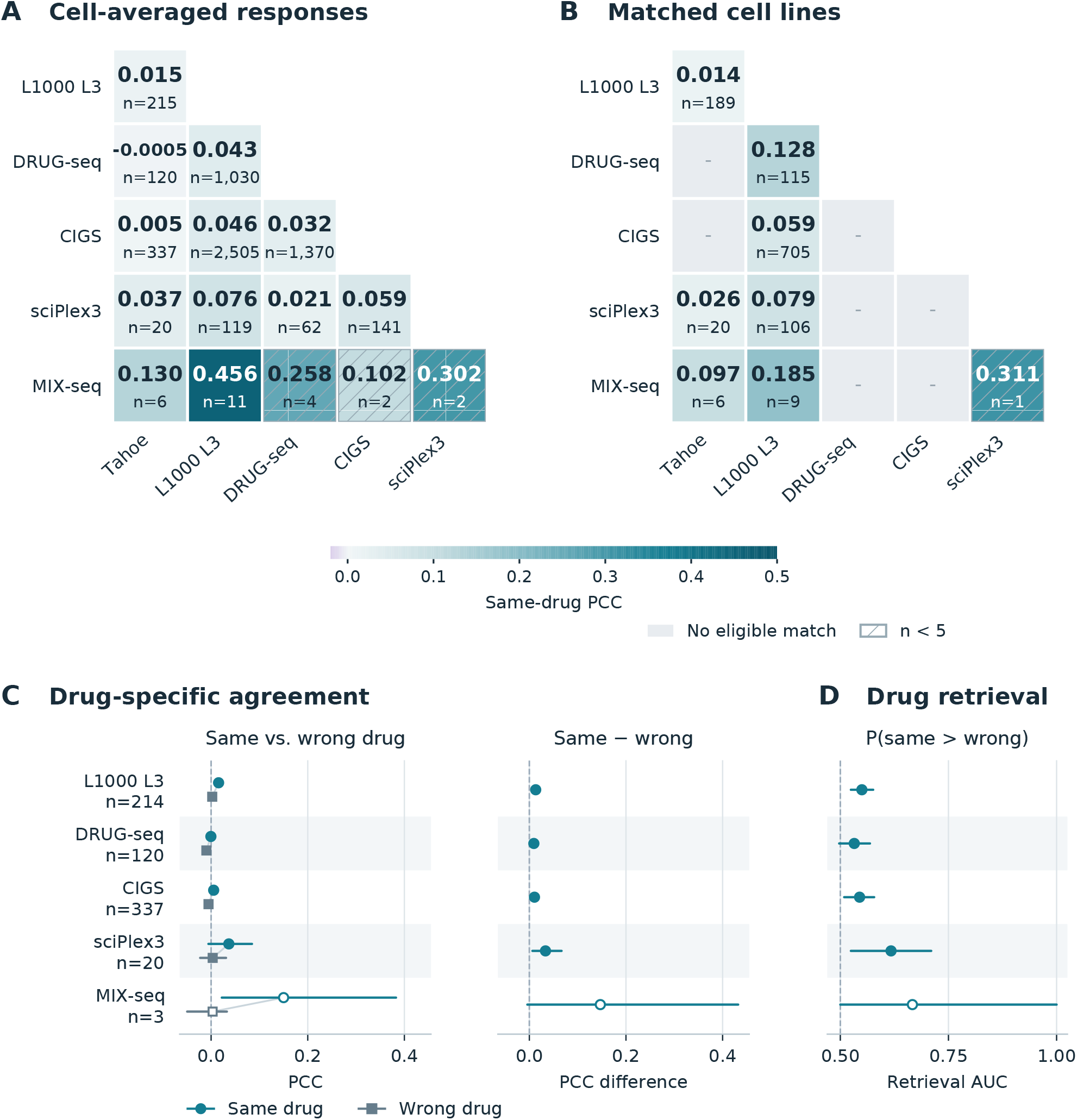
Cross-source response concordance and drug discrimination. A, concordance of same-drug responses averaged across cell lines within each source. B, same-drug concordance in matched cell lines. Eligible comparisons use 24 h treatment, doses within twofold, and 869 common genes. Tiles show mean PCC and the number of matched drugs; gray indicates no eligible match and hatching indicates fewer than five drugs. C, same-drug and background-matched wrong-drug PCCs, and their paired difference, for Tahoe queries against external sources. D, retrieval AUC: the probability that the same drug scores above an alternative, with chance at 0.5. C and D use the cell-averaged setting of A and candidate pools containing at least two drugs. Horizontal lines show 95% bootstrap intervals; open points indicate fewer than five query drugs.

### Drug main effects and interaction residuals dominate response variation in most datasets

To determine whether the measured variation primarily reflects differences among drugs or differences in how cells respond to them, we decomposed observed drug responses into an overall term, cell effects, drug main effects, and interaction residuals (Figure 3). For a complete drug-by-cell panel, the overall term is the mean expression change across drugs and cell lines. Cell effects describe each cell line’s average departure from this overall response across drugs, whereas drug main effects, denoted β, describe each drug’s average departure across cell lines. The interaction residual, denoted γ, captures how a particular drug-cell combination departs from the response expected by adding the overall, cell, and drug effects. It therefore represents combination-specific variation after accounting for both average patterns.

We analyzed all eligible observations in each retained dataset release on both the native measured gene axis and the shared gene set. Contributions were expressed as shares of the weighted squared response. In incomplete panels, additive fits and order-averaged allocation of cell and drug contributions accounted for the observed responses without imputing missing entries.

Drug main effects and interaction residuals together accounted for most response variation in Tahoe, L1000 L3, CIGS, and sciPlex3. Overall and cell-average responses alone therefore did not describe the dominant variation in these datasets. The balance differed among resources: L1000 L3 and CIGS had substantial drug main effects, whereas MIX-seq was dominated by cell effects. Because the resources profiled different sets of drugs and cell lines, these proportions describe their observed experimental panels rather than intrinsic biological properties of the measurement platforms.

Gene coverage also changed the apparent composition of the response. On Tahoe’s native gene axis, drug main effects accounted for approximately 67% of the squared response and interaction residuals for 15%; on the shared gene set, the corresponding shares were approximately 40% and 45%. L1000 L3 and CIGS showed smaller changes between gene sets. The relative contributions of drug main effects and interaction residuals are therefore not fixed properties of a dataset independent of the genes analyzed. Shared genes provide a common evaluation space, while native genes retain each resource’s broader measurement coverage.

Experimental design also determines which components can be distinguished. DRUG-seq profiled U2OS alone, allowing variation among drugs to be described within that cell line but preventing separation of a drug effect shared across cell lines from a U2OS-specific interaction. Figure 3 therefore displays this contribution separately as between-drug variation in a single cell line.

Drug main effects and interaction residuals comprised the major response components in most multicell datasets, but their relative contributions depended on gene coverage and experimental design. Interaction residuals also contain measurement error and other unexplained experimental variation. A large residual share therefore does not, by itself, establish stronger biological interactions or quantify the proportion of learnable signal.

**Figure 3.**
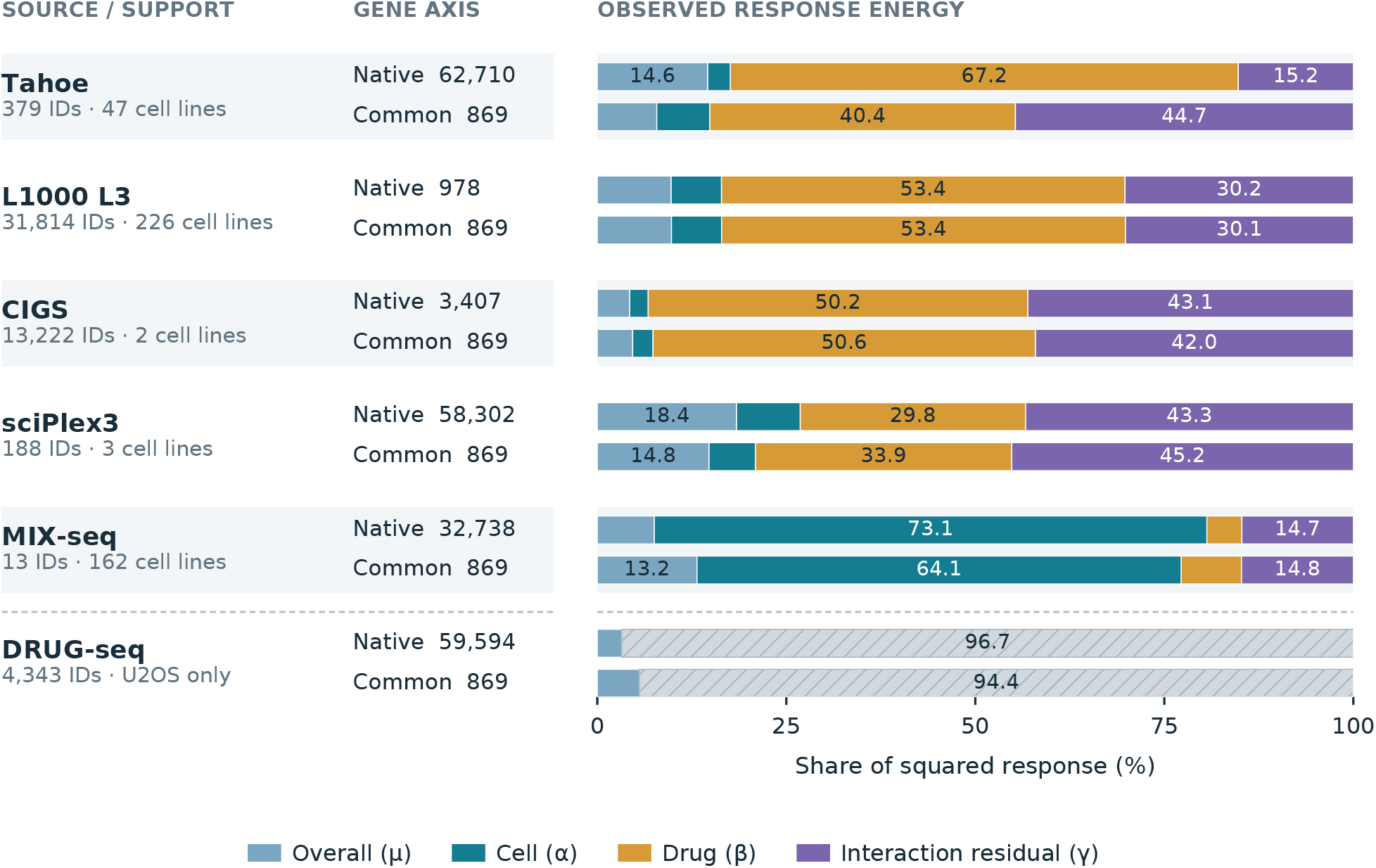
Composition of drug responses within each dataset. All eligible observations within each source are decomposed on native measured genes and 869 common genes. Colors indicate the grand response, cell effect, drug main effect, and interaction residual; numbers give their percentage contributions to the observed response sum of squares. Hatched DRUG-seq variation represents between-drug differences in U2OS alone, which cannot be separated into a cross-cell drug main effect and interaction. Interaction residuals include measurement error and other unexplained variation.

### Drug-specific main effects are a key bottleneck in the prediction framework examined

To determine which response components could be recovered from control expression, we examined a fixed Tahoe baseline that received no drug identity or molecular structure information. The model used matched control-cell expression as its sole input and predicted the complete drug-induced expression change through three hidden layers of 2,048 units each. We then applied the same decomposition to predicted and measured responses to evaluate their overall, cell, drug, and interaction components (Figure 4A). These components were derived from one complete prediction, rather than from four separately trained models.

On the established five-fold development panel of 153 drugs, the control-expression baseline achieved a complete-response PCC of 0.582, but recovery differed substantially among components. Correlations were higher for the overall and cell effects and lower for drug main effects and interaction residuals (Figure 4B). The latter two components together accounted for 95.9% of complete-response mean squared error (MSE; Figure 4C). An appreciable overall prediction correlation could therefore coexist with substantial unresolved drug-related variation.

This pattern was consistent with the model’s input restriction. Different drugs associated with exactly the same control-expression input necessarily received identical predictions, preventing the model from distinguishing their responses. We isolated the drug main-effect and interaction components that varied among drugs sharing this input and found that they accounted for 78.2% of total prediction error (Data S2). The baseline’s principal information gap was therefore concentrated in differences among drugs exposed to the same control-expression background.

We next asked whether a drug’s average response across cells contained information about its cell-specific interactions. We learned a mapping from β to γ using only the corresponding training data and supplied the measured β of evaluation drugs as input to predict the interaction component not represented by the baseline. This analysis tested the predictive relationship between drug main effects and interactions.

On the development panel, adding only the inferred interaction correction, without adding measured β itself, increased PCC and reduced MSE for both the overall interaction component and the complete response. Complete-response PCC increased from 0.582 to 0.612 (Figure 4D). Measured drug main effects thus contained interaction-predictive information that the control-expression baseline could not access. Because the input was derived from measurements of the evaluation drug, this was a measured-input diagnostic rather than a prediction of a previously unmeasured drug.

To test whether this relationship extended to independent experimental measurements, we used β measured on one plate to predict γ on another. After subtracting each plate’s own controls and removing its shared additive components, cross-plate interaction prediction remained positively correlated with the measured target and exceeded three training-input permutation controls (Figure 4E). On the same support of 47 shared drugs, mean interaction-prediction PCC decreased from 0.713 within plates to 0.387 across plates. The predictive relationship therefore persisted across the tested plate pair and could not be explained solely by additive variation shared within a single plate.

We also explored how drug main effects might be obtained for unmeasured drugs, using molecular structure, target information, and external same-drug measurements. Figure S2 summarizes ten core configurations with verified results. Structure-based representations and predicted targets generally yielded weak predictive correlations. Some annotated-target and CIGS response pipelines showed positive contrasts against their training-label permutation controls, but correlations remained low, and the pipelines differed in drug coverage, model form, and holdout rules. The complete exploratory inventory and development evaluations are retained in Data S2. These results indicate that some available inputs contain drug-specific information, but they have not yet established consistently accurate main-effect prediction.

Together, these findings identify accurate prediction of drug-specific main effects as a key bottleneck in the framework examined here. Measured β contains information about both the average drug response and its cell-specific interactions, and the latter relationship persists across independently measured plates. Improving β prediction for unmeasured drugs is therefore a promising route toward recovering both drug main effects and interaction responses.

**Figure 4.**
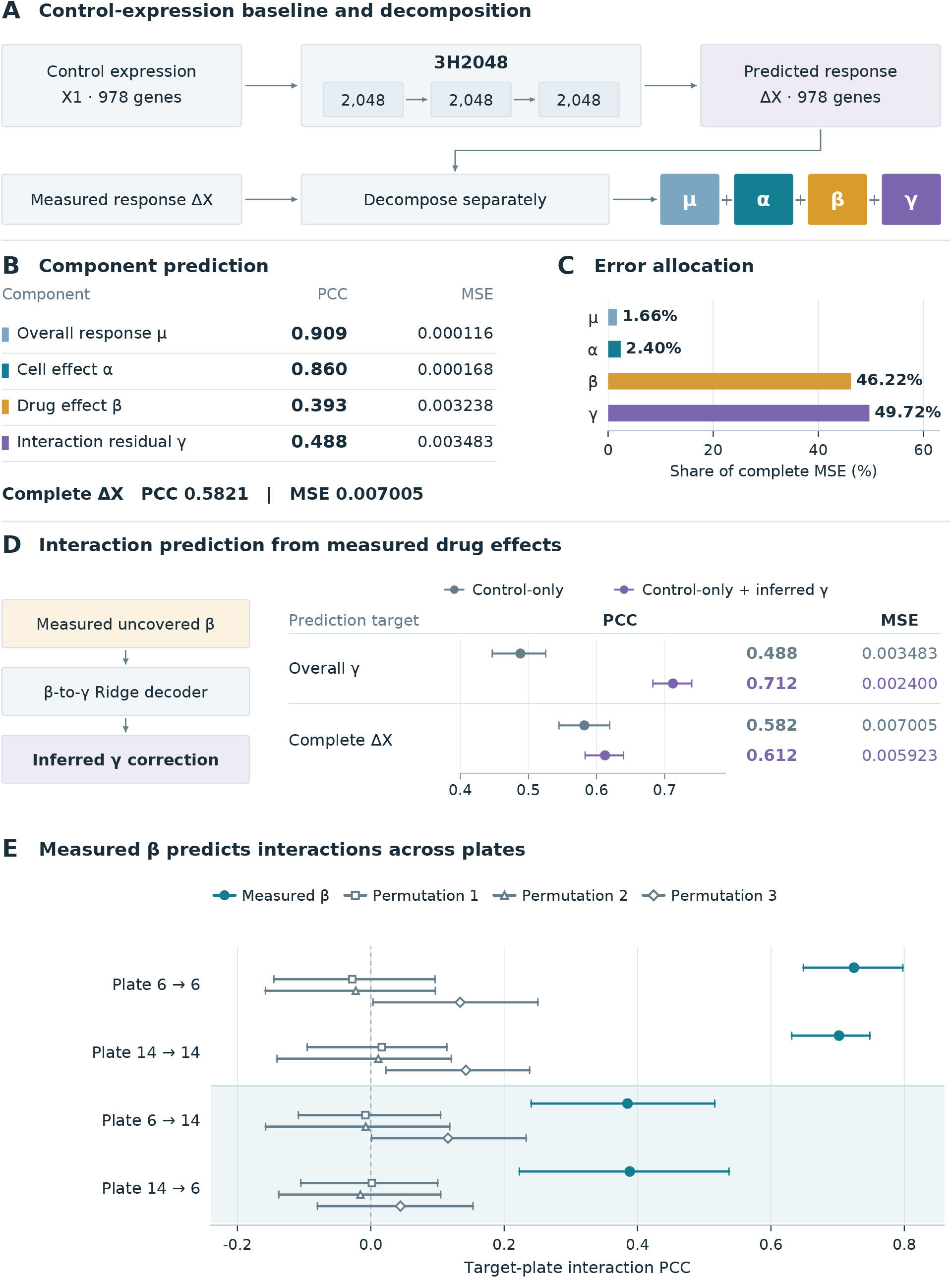
Control-expression baseline and interaction information in drug main effects. A, prediction from control expression alone and response decomposition. The model predicts the complete drug response; predictions and measurements are decomposed identically. B, component PCC and MSE. C, component contributions to complete prediction MSE. D, measured drug main effects (β) are passed through the training-fitted G5 decoder to infer an interaction correction not captured by the baseline. Only the inferred interaction is added; measured β itself is not added. Right, overall interaction and complete-response predictions before and after correction. Gray denotes the baseline and purple denotes the corrected prediction. E, measured source-plate β predicts target-plate interaction. Arrows indicate source-to-target plate direction, shading marks cross-plate predictions, and three training-input permutation controls are shown. B-D use 153 drugs and E uses 47 drugs shared by both plates; both settings use 47 cell lines, 978 genes, and 5 μM. Horizontal lines show 95% bootstrap intervals. D and E are measured-input diagnostics.

## Discussion

Existing drug-perturbation transcriptomic resources have broadened the range of drugs and cellular backgrounds that can be studied. We found that treated expression could remain highly consistent between independent repeats, whereas drug-induced expression changes were substantially less repeatable. Stable cellular expression backgrounds can therefore obscure variability in drug responses. For studies seeking to learn or validate drug effects, these changes are the measurements that need to be reliable. High correlation between treated expression profiles alone can overstate the reliability of the drug-response information available for such analyses.

Cross-source comparisons revealed a further challenge. The same drug often elicited poorly concordant responses across datasets, even when cell identity, treatment duration, and dose were closely matched. These discrepancies limit the use of one source to corroborate findings in another or to serve as a proxy for its measured responses.

Data integration therefore requires addressing differences in measurement itself; harmonizing gene identifiers and experimental annotations is only a starting point. Nevertheless, some sources retained a same-drug matching advantage, indicating that shared drug-specific information was not entirely absent. This information may help identify related response patterns or provide predictive priors, but its value needs to be assessed for the intended task.

Response decomposition clarified the information missing from the prediction framework. The control-expression baseline could use cellular state and experimental background, but it could not distinguish drugs associated with identical inputs. An appreciable complete-response correlation therefore did not establish that the model had recovered differences among drug responses. Evaluating components separately identifies which predictive abilities support the overall score and whether additional drug information improves responses that the baseline could not distinguish.

Drug main effects (β) have a dual role in this framework: they describe differences among drugs in their cell-averaged responses and contain information predictive of cell-specific interactions. This relationship persisted across independently measured plates, providing a basis for jointly modeling average drug effects and interaction responses. The unresolved challenge is obtaining accurate drug main effects for drugs not yet profiled in the target system. Explorations using molecular structure, target information, and external response representations identified positive signals in a small number of pipelines, but main-effect prediction remained weak overall. Accurate prediction of drug main effects is therefore a key bottleneck in the framework examined here and a promising direction for improving cell-specific response prediction.

Future data resources would benefit from shared reference measurements that permit direct comparison across sources: the same drugs and cell lines, closely aligned doses and treatment durations, and independent controls and repeats within each source. Such measurements would help establish whether a drug response can be reproduced across experimental settings. For modeling, complete-response and component-level evaluations should complement one another. Complete-response metrics assess whether final predictions improve, whereas component-level analyses clarify where those improvements arise. Together, they can establish whether additional drug information addresses the identified prediction gaps and make methodological improvements easier to interpret and compare.

The comparisons in this study remain constrained by differences in drug and experimental-condition coverage, which prevent isolation of the contributions of measurement noise and individual experimental factors. The prediction analyses used a fixed baseline and an established development panel. Cross-plate validation covered one plate pair and tested the interaction-prediction potential available when drug main effects were measured. Complete-response gains varied with transfer direction. For drugs not yet profiled in the target system, the next question is how much interaction-predictive information is retained when measured drug main effects are replaced by model predictions, and whether that information supports consistent improvements in complete-response prediction.

## Acknowledgments and funding

Computing facilities were provided by the University of Cape Town ICTS High Performance Computing team (https://doi.org/10.5281/zenodo.10021612).

Sharon Prince received funding from the University of Cape Town (UCT Vision 2030 Grand Challenges Programme); the National Research Foundation of South Africa (Competitive Programme for Rated Researchers, SRUG190306422357 and SRUG2204224267); the International Centre for Genetic Engineering and Biotechnology (Collaborative Research Programme, CRP/ZAF20-01); the Poliomyelitis Research Foundation; and the South African Medical Research Council (Self-Initiated Research Grant). The views expressed are the authors’ and do not necessarily reflect those of SAMRC or other funders.

## Author contributions

J.B., S.P., and G.S.N.: Conceptualization. J.B.: Methodology, Software, Formal analysis, Investigation, Data curation, Visualization, and Writing - original draft. S.P. and G.S.N.: Supervision, Funding acquisition, and Writing - review and editing.

## STAR Methods

### Key resources table

| Resource | Source | Identifier |
|---|---|---|
| Tahoe-100M expression and annotations | Zhang et al. [1] | 10.1016/j.cell.2026.08.035 |
| L1000 Level 3 expression and RNA-well annotations | Subramanian et al. [2] | 10.1016/j.cell.2017.10.049 |
| DRUG-seq counts and plate annotations | Ye et al. [3] | 10.1038/s41467-018-06500-x |
| sciPlex3 counts and replicate annotations | Srivatsan et al. [4] | 10.1126/science.aax6234 |
| CIGS counts and compound catalog | Xiang et al. [5] | 10.1038/s41592-025-02781-5 |
| MIX-seq counts and cell assignments | McFarland et al. [6] | 10.1038/s41467-020-17440-w |
| KPGT molecular features | Li et al. [9] | 10.1038/s41467-023-43214-1 |
| Control-expression baseline and G5 decoder | This study | DrugPerturbDB: data/config/models.json |
| Analysis code, source tables, and CPU environment | This study | https://github.com/baijinming97/DrugPerturbDB |

File-level download URLs, sizes, SHA256 checksums, gene mappings, and fixed data splits are supplied with the repository. The CPU environment specifies Python and package versions, CPU PyTorch, and the R dependencies needed for DRUG-seq preparation.

### Resource availability

#### Correspondence

Further information and requests should be directed to Sharon Prince or Geoff S. Nitschke.

#### Materials availability

This study did not generate new biological materials.

#### Data and code availability

Source data are available from the studies listed above. Analysis code, numerical results, identity and condition annotations, gene mappings, and ten frozen models are available in DrugPerturbDB. The repository provides download, preparation, and analysis commands with a locked CPU environment. Version-matched CIGS count inputs and fixed feature matrices for the exploratory comparisons are included. Predictive analyses use the supplied frozen weights. Code and study-generated materials are distributed under CC BY-NC 4.0; external resources retain their original terms.

#### Experimental model and study participant details

We reanalyzed six drug-perturbation datasets from human cell lines [1–6], without new participant, animal, or cell-culture experiments. Source-wide analyses included all observations meeting annotation, control, and quality requirements in the specified releases, without first intersecting drugs with Tahoe. Cross-source analyses additionally required matched identities and treatment conditions.

Tahoe descriptions covered 379 drug IDs, 47 cell lines, and 52,886 qualified conditions at 0.05, 0.5, and 5 μM. Predictive analyses retained the fixed five-fold chemical-group protocol: a 261-drug candidate panel and 153 unique development-VALID drugs. Descriptive inclusion was independent of model roles. The model TEST partition was not evaluated, and expanded descriptions were not used to retrain or select models.

### Method details

#### Identity, gene axes, and response construction

Drug matching used source structures, official compound identifiers, CAS and PubChem information, and parent-structure reconciliation preserving stereochemistry. Conflicting or unresolved identities were not forcibly merged. Cell identities were reconciled to Cellosaurus; unresolved assignments were excluded from same-cell comparisons. Response similarity was not used for matching.

Each source was analyzed on native measured genes and a fixed set of 869 common Entrez genes. The latter retained the eight-source intersection defined during analysis design. The 882-gene intersection of the six current sources was evaluated separately on identical matched samples. Missing or ambiguous gene mappings were masked, not assigned zero. L1000’s native axis contained 978 measured landmark genes, excluding inferred expression. Duplicate sciPlex3 Ensembl entries were consolidated and nonhuman genes excluded from its native human axis.

Count-based sources were normalized by the original full-library total before gene subsetting and transformed as log2(CPM + 1); log2(CPM + 100) was the processing sensitivity. Tahoe retained author-provided linear normalized expression, and L1000 L3 retained source normalization. Responses were treated expression minus a matched control from the same cell and experimental batch.

Tahoe plate-level pseudobulks required at least 50 full-QC cells and retained the source-qualified 47-cell panel. Descriptive condition profiles pooled eligible repeat plates by full-QC cell count; reconstruction of the frozen predictor instead retained its original equal-plate aggregation. sciPlex3 and MIX-seq pseudobulks were defined by cell, drug, dose, time, and independent replicate, required at least 20 cells, and required a corresponding control. DRUG-seq used nonempty DMSO wells on the same physical plate and excluded zero-library wells. L1000 controls were matched within RNA plate and condition; technical rereads of the same physical RNA well were merged.

#### Independent repeatability

Replicate pairs matched drug, cell, dose, and time but came from distinct physical plates or independent biological replicate units with nonshared control pools. Same-plate pairs sharing controls were excluded. Tahoe independence was verified at the physical-plate control-pool level.

Across-gene Pearson correlations were computed separately for normalized treated expression and treated-minus-control responses, then averaged across repeat pairs within condition and across conditions within drug. External sources used equal drug weight; Tahoe used equal chemical-group weight and equal drug weight within group. Within each endpoint, native and common gene analyses used the same valid pairs: a pair with undefined PCC on either axis was excluded from both. This rule was applied separately to expression and response. Repeatability was not used for attenuation correction or converted into a prediction ceiling.

#### Cross-source agreement and same-drug retrieval

Primary comparisons used 24 h treatments and all positive-dose pairs differing by no more than twofold. Dose-pair estimates were averaged within drug and cell before cell and drug aggregation. Responses averaged across each source’s available cells and responses in confirmed matching cells were reported separately. Comparisons involving Tahoe used equal chemical-group weight, with equal drug weight within group; other source pairs used equal drug weight.

Same-drug and wrong-drug scores used candidate pools sharing treatment time and the two source-dose settings, with at least two drugs. Retrieval AUC was the mean of I(same > wrong) + 0.5 × I(same = wrong) over alternative drugs. Matched-minus-mismatched PCC and retrieval AUC used identical queries.

For the six source pairs with independent repeats on both sides, within-source and cross-source correlations were computed on identical drugs, cells, doses, and 869 genes. All estimates used single-repeat profiles, averaging eligible combinations within condition. Source pairs without joint repeat support were not scored in this comparison. Sensitivities included 882 genes, exact doses, the larger pseudocount, equal-drug weighting, and plate deletion or repeat averaging where feasible.

#### Response decomposition

Complete weighted panels were decomposed for each gene as Y(c,d) = μ + α(c) + β(d) + γ(c,d). The grand mean μ, cell deviation α, and drug deviation β were obtained from weighted marginal means; γ was the remaining interaction residual.[8] Measurements and predictions were decomposed separately using identical weights.

In Tahoe, drugs were grouped by their complete 47-cell control-expression pattern. Subtracting the weighted drug mean within these groups defined the drug main-effect component not represented by the baseline. The corresponding interaction component was obtained by constrained projection, with zero weighted mean across cells for each drug and across drugs within each cell-context group. Predictive corrections fixed contexts containing fewer than three drugs to zero; descriptive decomposition used the full constraint space. Projection therefore depends on reference drugs in the evaluation panel.

Source-wide component energies assigned equal total weight to each drug, shared equally among its observed cell, dose, time, and library conditions. Incomplete panels were stratified by dose, time, and library and fitted by weighted least squares using observed entries only. Cell and drug contributions were averaged over their two orders of entry; stratum energies were combined using their weight mass. These contributions and the residual sum to the observed response sum of squares. Designs unable to distinguish drug and interaction effects, including single-cell-line panels, were marked separately. The decomposition describes observed variation, including measurement error.

#### Fixed baseline and exploratory predictions

The frozen control-expression baseline was a seed1337 multilayer perceptron with three 2,048-unit hidden layers, ReLU activations, dropout 0.1, and 978 inputs and outputs. Matched control expression X1 predicted complete ΔX without drug identity or structure. Its original loss was 0.5 × MSE(X1 + predicted ΔX, X2) + 0.5 × (1 - condition-wise ΔX PCC). Checkpoints had been selected on outer VALID; these weights remained fixed. Component scores were derived from the same complete prediction, not from separately trained branches.

Figure S2 reports ten fixed configurations under chemical-group holdout or joint chemical-group and known-mechanism holdout. Inputs comprised molecular structure, target information, and external measurements; the target was the drug main-effect component not represented by the baseline. Inputs, model forms, response ranks, and drug support are specified in Data S2 and the supplied configurations. Annotated-target priors used target-linked L1000 genetic-perturbation signatures, distinct from same-drug chemical-treatment signatures.

Preprocessing, target bases, hyperparameters, and shrinkage were determined within the corresponding TRAIN fit blocks. The Ridge/three-program configurations used up to 32 input PCs, four chemical-group holdout blocks, and three-fold inner selection of the regularization parameter from {0.1, 1, 10, 100}. Each configuration was compared with three whole-drug training-label permutations (seeds 1337, 2026, and 3407), exchanging labels between equally sized chemical groups; held-out inputs remained unpermuted. Primary exploratory scores used the outer-TRAIN context-centered target; held-block centering was reported separately. Missing inputs retained the baseline in complete-response comparisons.

#### Drug-main-effect-to-interaction mapping

The G5 decoder used measured drug main effects to predict interaction responses not represented by the baseline. Selection and out-of-fold shrinkage used four complete control-context blocks, excluding fitting drugs whose chemical groups overlapped the held-out block. Hyperparameter selection, when needed, was nested within those blocks.

Projected 5 μM TRAIN interactions were arranged as drug-by-(cell × gene) matrices. A chemical-group-weighted rank-20 basis was fitted by iterative low-rank reconstruction of missing eligible entries, with at most 250 iterations and a relative objective-change tolerance of 1e-5. Drug main effects were centered within control-context groups; PCA used the unweighted TRAIN mean and up to 32 components, standardized by TRAIN score standard deviations. Available 0.5 and 0.05 μM TRAIN panels used the same basis and preprocessing, with dose covariate 1 + log10(dose/5). Each dose block’s weights were normalized before pooling for Ridge fitting. Predicted interactions were constrained and shrunk using TRAIN calibration before addition to the baseline; measured drug main effects themselves were not added.

Cross-plate validation fixed Ridge regularization at 100 and the correction coefficient at 1, without a new search. Evaluation drugs and their chemical groups were excluded at every training dose. All four within- and cross-plate directions for plates 6 and 14 used the same 47 drugs, 47 cell lines, 978 genes, and 5 μM conditions. Source-plate drug means used all 47 cells.

Three controls permuted TRAIN inputs by whole drug within joint dose-coverage patterns, using a shared mapping across doses (seeds 20261103, 20261104, and 20261105). Only the input-to-code Ridge map was refitted; basis and preprocessing remained fixed, and evaluation inputs were unpermuted. Both transfer directions were retained and weighted equally when summarized.

### Quantification and statistical analysis

PCC was computed across genes within the specified condition or component and aggregated with the stated support weights; MSE used the same weights. Predictive analyses used equal chemical-group weight, equal drug weight within group, and equal cell and gene weights. Repeated VALID roles shared each drug’s weight.

The 95% intervals were the 2.5th and 97.5th percentiles of 2,000 paired bootstrap draws. External source-wide and external-to-external comparisons resampled drug IDs; Tahoe and comparisons involving Tahoe resampled chemical groups. Model and control contrasts used the same draws. Cross-plate analyses re-estimated centering and projection in each draw while holding models fixed. Single-drug estimates had no drug-level interval; cross-source intervals required at least three resampling units. Panels with fewer than five drugs were marked descriptive.

For complete balanced panels, component MSEs were checked to sum to complete MSE. Pure-interaction corrections were required to preserve μ, α, and β and to produce equal changes in complete and interaction MSE. Undefined measurement and complete-response correlations were omitted rather than filled with zero. The retained increment-score convention assigned zero to degenerate corrections; the single-plate baseline’s undefined interaction PCC was not treated as zero.

Intervals describe uncertainty conditional on the fixed samples and analysis, not historical model search or retraining. The three permutations are fixed negative controls rather than a permutation-test null distribution. Cross-plate intervals condition on the tested plate pair. Reported analyses can be recomputed in the locked CPU environment; independent checks cover representative repeat correlations, cross-plate prediction, sample and gene axes, role isolation, and energy closure. Numerical results, configurations, and pairing lists are supplied in Data S1, Data S2, and the repository.

## Declaration of generative AI assistance

OpenAI Codex (OpenAI; desktop application version 26.1103.3517.0 at final preparation) assisted with code development, documentation, manuscript preparation, language editing and figure-code preparation. Numerical analyses, statistical figures and schematics were generated by executable Python code; statistical values came from measured data and were not generated or altered by an image model. The authors are responsible for the scientific content and its verification.

