## Supplementary Figures for "Reproducibility and cross-dataset concordance of drug-induced transcriptional responses"

### Supplemental figures

#### Figure S1. Effects of processing and matched support on measurement comparisons

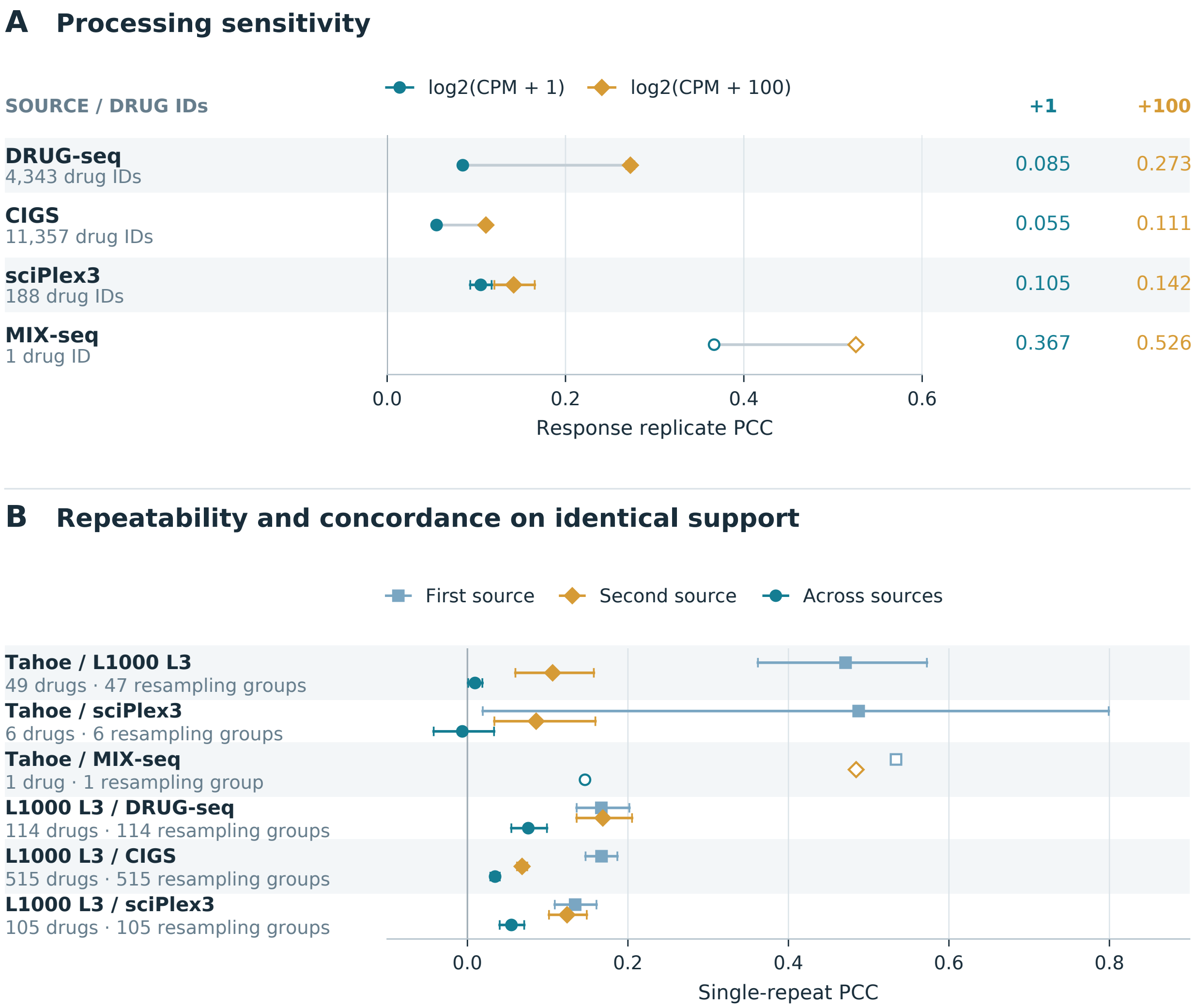


A, response repeatability after log2(CPM + 1) or log2(CPM + 100), using identical replicate pairs and 869 common genes. Gray lines connect estimates from the same source.

B, within-source repeatability and cross-source concordance on shared conditions with independent repeats in both sources, for all six eligible source pairs. First and second source follow row-label order; all three comparisons use identical matched conditions and genes. Horizontal lines show 95% bootstrap intervals; open points indicate fewer than five drugs. Single-drug comparisons have no drug-level interval.

#### Figure S2. Exploratory prediction of drug-specific main effects

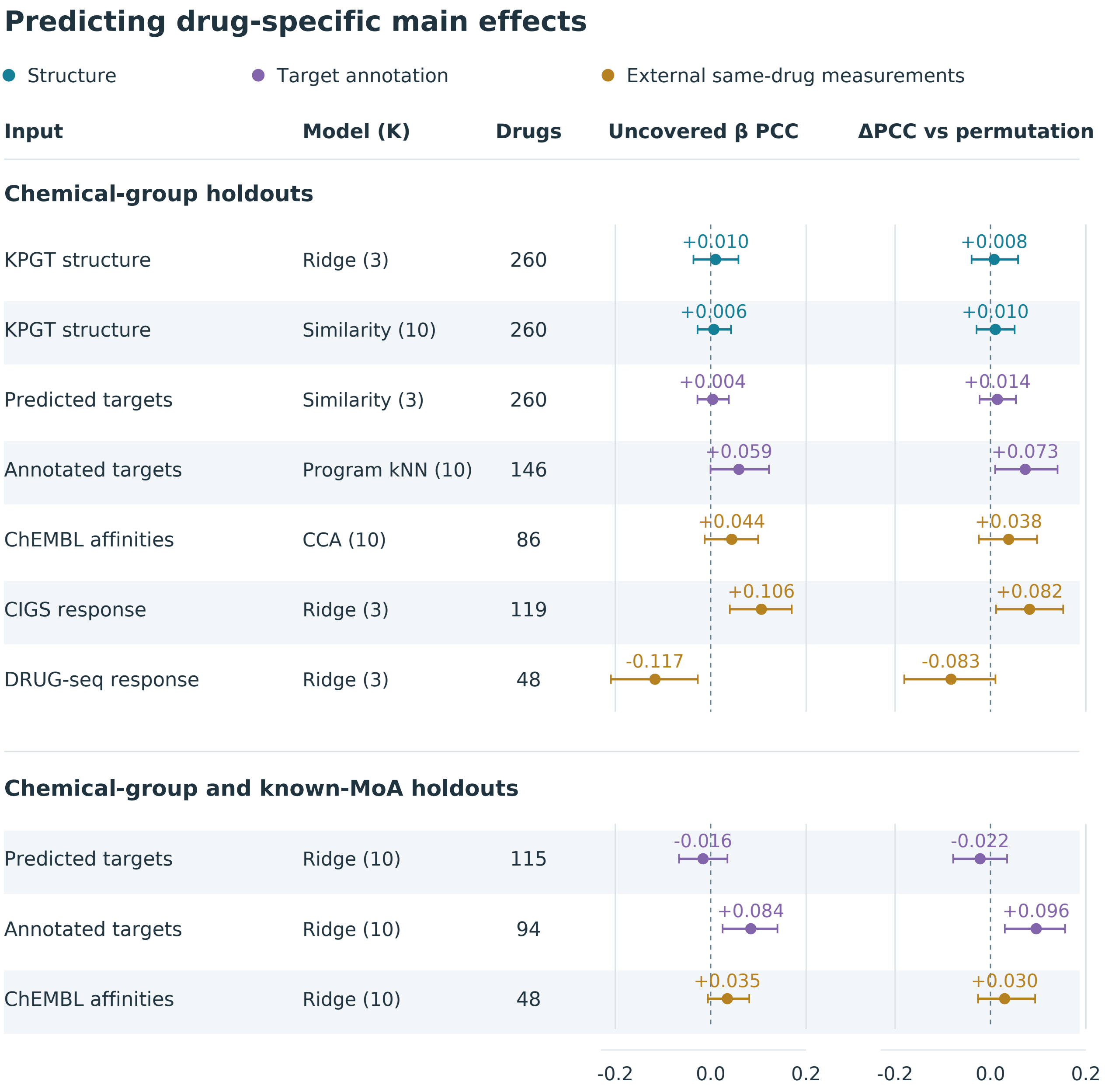


Ten configurations predict the drug main-effect component not captured by the baseline. Upper, chemical-group holdout; lower, chemical-group and known mechanism-of-action holdout. Left, weighted mean drug-wise correlation across 978 genes. Right, paired correlation difference relative to three training-label permutation controls. Horizontal lines show 95% bootstrap intervals.

K denotes the number of response programs; Similarity, similarity-weighted prediction; CCA, canonical correlation analysis; and MoA, mechanism of action. Annotated targets uses target-linked L1000 genetic-perturbation signature priors. All configurations are exploratory holdout evaluations within the training data.
